# Alström syndrome proteins are novel regulators of centriolar cartwheel assembly and centrosome homeostasis in *Drosophila*

**DOI:** 10.1101/2023.09.13.557518

**Authors:** Marine Brunet, Joëlle Thomas, Jean-André Lapart, Maria Giovanna Riparbelli, Giuliano Callaini, Bénédicte Durand, Véronique Morel

**Affiliations:** Université Claude BERNARD Lyon 1, Lyon, France; MeLiS – CNRS-UMR5284; INSERM-U1314; Università degli Studi di Siena, Siena, Italy

**Keywords:** #centriole duplication, #cartwheel, #Ana2, #Plk4, #Alms1, #Alström syndrome

## Abstract

Centrioles play central functions in cell division and differentiation. Alterations of centriole number or function lead to various diseases including cancer or microcephaly. Centriole duplication is a highly conserved mechanism among eukaryotes during which Ana2/STIL phosphorylation by Plk4 allows the recruitment of the cartwheel protein Sas-6 to initiate procentriole assembly. Here we show that the two *Drosophila* Alms1 proteins (Alms1a and Alms1b), orthologs of the human ALMS1 protein associated with the Alström syndrome, are novel regulators of the assembly of the pro-centriole cartwheel and pericentriolar material (PCM). Using Ultrastructure Expansion Microscopy we show that Alms1a is a PCM protein loaded at the proximal end of centrioles at the onset of their assembly in several fly tissues, while Alms1b caps the proximal end of only maturating centrioles. We demonstrate that chronic loss of Alms1 proteins affects PCM maturation, whereas their acute loss impairs the onset of procentriole formation by reducing Ana2 amplification leading to complete failure of Sas-6 recruitment. Our work identifies Alms1 proteins as novel centriolar components that finely tune centrosome homeostasis and the initiation of cartwheel formation in *Drosophila*. It also reveals strong buffering capacity of tissues in response to perturbations of the centriole assembly process.

## Introduction

Centrioles are highly conserved, small, cylindrical organelles composed of nine triplets of microtubules. They are surrounded by a complex array of proteins, the pericentriolar material (PCM), and lie at the core of the centrosomes that serve as microtubule-organising centres (MTOC), playing a central role in cell division. Alterations of centriole number are associated with numerous pathologies including cancers ^1^. Centriole duplication must therefore be tightly regulated and coordinated with the cell cycle ^2^.

Non dividing cells normally contain a pair of connected centrioles, an older, the mother, and a younger, the daughter, arranged perpendicularly. As the cell enters S-phase, centrioles disconnect in a process called disengagement and initiate their duplication. A procentriole forms on the side of each centriole, grows to become a daughter centriole and recruits PCM. The two duplicated centrosomes hence formed organise the mitotic spindle and, ultimately as cells divide, each daughter cell inherits a single centrosome made of a mother and a daughter centriole.

Centriole duplication relies on a few sets of proteins highly conserved across species and which functions are highly hierarchised. Centriole duplication initiates with the recruitment of the kinase PLK4/Plk4 (or Sak) by the PCM proteins Cep152/Asl and Cep192/Spd2 at the proximal side of the centriole ^3–5^. Plk4 initially forms a cylinder around the centriole and next resolves to a single dot which corresponds to the future site of procentriole assembly ^6,7^. This reorganisation relies on a complex cascade of trans autophosphorylations ^8–10^ leading to Plk4 activation. This initiates the sequential phosphorylation on multiple sites of STIL/Ana2 which can then recruit SAS-6/Sas-6 ^6,11,12^ to form the cylindrical nine-fold symmetry array of the cartwheel. The latter serves as a template for Sas-4-controlled deposition of microtubule triplets initiating procentriole elongation ^13,14^.

ALMS1 is a centrosome and cilia associated protein characterised by a C-terminal ALMS domain involved in the localisation of the protein at centrioles and basal bodies ^15,16^. Mutations in *Alms1* are responsible for a rare ciliopathy, the Alström syndrome ^17,18^, but how ALMS1 controls ciliary functions is still enigmatic. Two *Alms1* genes have been identified in *Drosophila*, *alms1a* and *alms1b*, which share more than 80% sequence homology. Alms1a was shown to localise at the centrosome in asymmetrically dividing male germline stem-cells (GSC) and to be involved in centriole duplication in these cells, at the exclusion of other cell or division types ^19^.

We undertook to characterise Alms1a and Alms1b roles at centrioles. Using Ultrastructure Expansion Microscopy (U-ExM), we characterised Alms1a and Alms1b localisation with unprecedented resolution. We show that both localise at the proximal end of centrioles but with spatiotemporal differences suggesting specialised functions of each protein. We observe that acute depletion of both Alms1a and b by RNAi is associated with complete centriole duplication failure in various *Drosophila* tissues, while chronic loss of both proteins in a double KO mutant leads to centriole disengagement and reduced PCM recruitment, at least in the male germline. This shows that centriole duplication is a highly buffered process with strong compensatory mechanisms triggered in *alms1a,b* mutant situations. Finally, we place Alms1a,b in the molecular hierarchy of centriole duplication and show that they are involved in fine tuning Ana2 activity downstream of Plk4, Alms1a,b loss leading to complete failure of Sas-6 recruitment. Altogether our work demonstrates that Alms1a and b are novel players in the initiation of cartwheel formation and PCM assembly in *Drosophila*.

## Results

### Alms1a and Alms1b present different spatiotemporal localisations at centrioles

Alms1a has been proposed to play a critical role in the asymmetry between mother and daughter centrosomes in male germline stem-cells (GSCs) at the exclusion of any other cell type ^19^. To characterise the centriolar localisation of both proteins and their dynamics of recruitment during centriole duplication in different cell types (Fig. 1), we adapted the U-ExM protocol ^20^ to several *Drosophila* tissues (testes, larval brains and early embryos) expressing either Tomato tagged Alms1a or GFP tagged Alms1b under their native promoter in rescue conditions (in *alms1a* or *alms1b* mutant respectively, Fig. S1).

**Figure 1.**
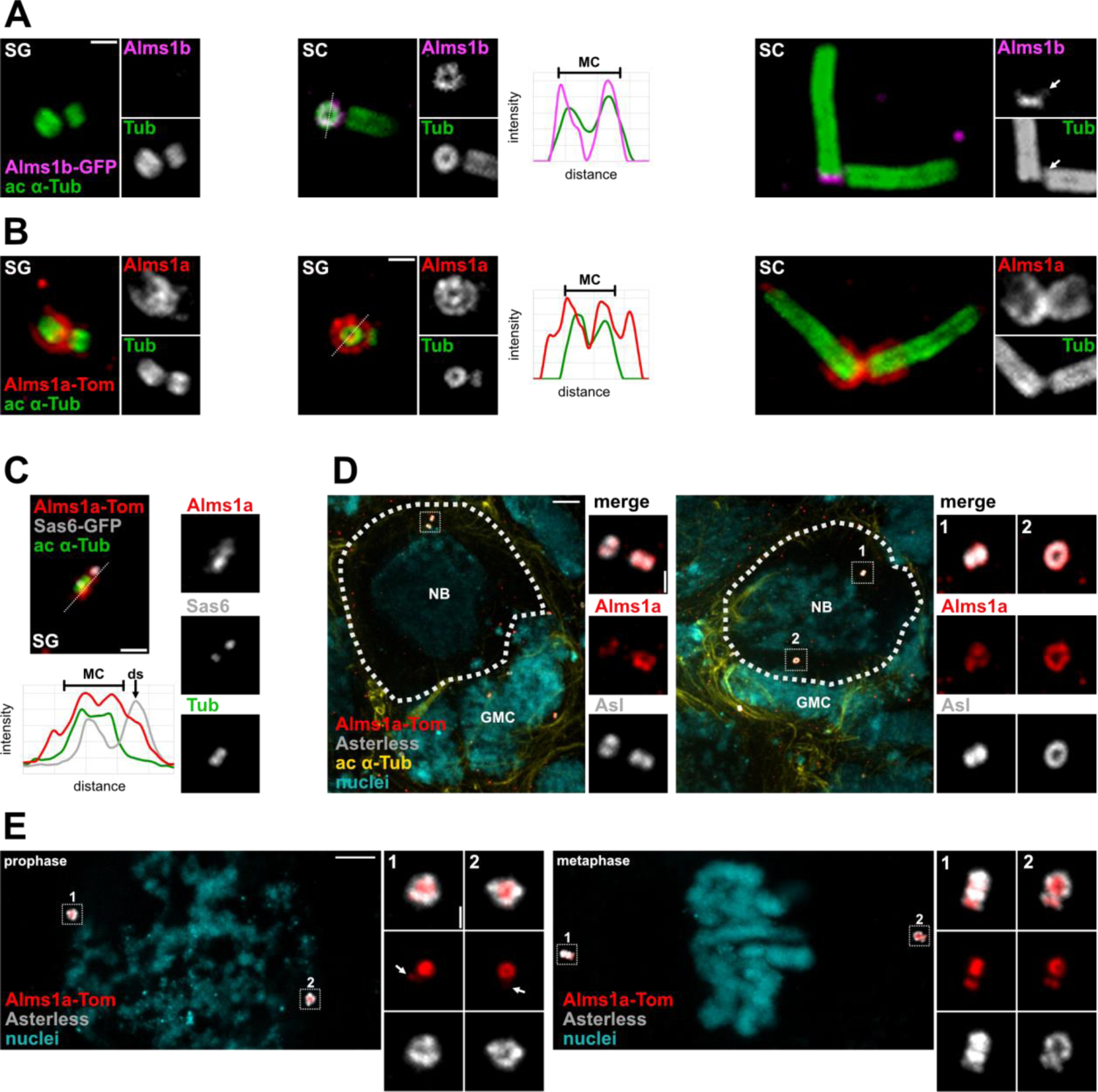
Alms1a and Alms1b show different spatial-temporal localisations. U-ExM images of : (A-C) Centriolar localisation of (A) Alms1b-GFP (magenta) and (B) Alms1a-Tomato (red) at early stages of spermatogenesis (spermatogonia SG; spermatocytes SC). (C) Relative localisation of Alms1a-Tomato (red) and Sas-6-GFP (grey) in a duplicating centriole. In all images, centriolar walls revealed with acetylated α-tubulin (green, ac a-tub), mother centrioles on the left, daughter centrioles on the right. Fluorescence intensity measures across mother and daughter centrioles (dotted line). MC bracket: mother centriole width. Arrows: duplication site (ds). Scale bars, 1 μm. (D) Alms1a-Tomato (red) localisation at centrioles in larval brain neuroblasts (NBs) and ganglion mother cells (GMCs). The left panel shows a NB in interphase. Insets: close up of the centriole pair, the mother centriole at the top and the daughter centriole at the bottom. Alms1a-Tomato forms a ring superposed to Asterless ring and is less abundant at the daughter than at the mother centriole. The right panel shows a dividing Nb. The two centrioles have separated and migrated on both sides of the cell. The bottom, “mother”, centriole will be inherited by the future GMC. Asterless (grey) labels the centriole, acetylated α-tubulin (yellow) the cytoskeleton and Hoechst (cyan) the nuclei. (E) Alms1a-Tomato localisation at centrioles in early embryos. Top panel : prophase. In the inset, the two centrioles have initiated their duplication as evidenced by the light Alms1a-Tomato concentration on the side of the centriole preceding the recruitment of the inner PCM protein Asterless. Bottom panel: metaphase with well advanced centriole duplication as shown in the insets by the accumulation of both Alms1a-Tomato and Asterless on the side of the mother centrioles. In these two stages, the cells are dividing, forming both asters and mitotic spindle. This explains the wide and intense Asterless staining with respect to Alms1a-Tom. This contrasts with NBs, GSCs and SCs were cells are not in the process of forming a spindle and in which Alms1a-Tom localisation is slightly wider than Asterless one. Asterless (grey) labels the centriole and Hoechst (cyan) the nuclei (cyan) (D-E) Scale bars, 5 or 1 μm (insets).

We first documented Alms1a-Tomato and Alms1b-GFP precise localisations during spermatogenesis (Fig. 1A-C). *Drosophila* spermatogenesis starts with the asymmetric division of a GSC which gives rise to a GSC and a goniablast (GB) that initiates differentiation into a spermatogonium (SG; Fig. S2A). SG undergo four symmetric divisions to generate a cyst of 16 cells, each containing two extremely small centrioles (also called minute centrioles). As SGs complete a pre-meiotic S phase and duplicate their centrioles, they differentiate into spermatocytes (SCs). In SCs, the 4 centrioles grow unusually long, reaching about 1.3 μm in length and all centrioles mature into basal bodies nucleating a primary-like-cilium ^21^. SCs finally undergo two meiotic divisions, forming 4 spermatids each (resulting in 64 spermatids for each initial GB) with one basal body from which the sperm flagellum emanates ^22^.

We observed that Alms1b-GFP is not detected at centrosomes in spermatogonia (Fig. 1A, SG). It is first recruited to the mother centriole in early spermatocytes, forming a ring which caps the proximal end of centriolar microtubules, while no Alms1b-GFP is observed on the daughter centriole (Fig. 1A, SC, middle). This strong asymmetry within the centrosome is maintained until the end of the spermatocyte stage, when faint Alms1b-GFP can be seen at the proximal side of the daughter centriole (Fig. 1A, SC, right, white arrow). Later, all spermatids exhibited equivalent amounts of proximal Alms1b-GFP (not shown), which suggests that its recruitment occurs rapidly at the very end of spermatocyte stage or during meiotic divisions.

In contrast, Alms1a-Tomato localises at centrosomes from the GSC stage onwards. Alms1a-Tomato forms a ring at the base of the centriole wall and a sleeve surrounding it (Fig. 1B; Fig. 2A,B). It is loaded during each centriole duplication event at the onset of cartwheel assembly (Fig. 1B, SG; 1C). Alms1a-Tomato is indeed found surrounding the cartwheel component Sas-6-GFP at the centriole duplication site, before the observation of procentriole assembly based on acetylated α-tubulin staining (Fig. 1C). Alms1a-Tomato then accumulates on the daughter centriole and at mid-late spermatocyte stages, both mother and daughter centrioles present similar amounts of Alms1a-Tomato at their proximal end (Fig. 1B, SC). Together, these results indicate that Alms1a is associated with each centriolar duplication event in spermatogenesis, from stem-cells to spermatogonia, whereas Alsm1b is associated with post-duplication maturation of the centrioles.

**Figure 2.**
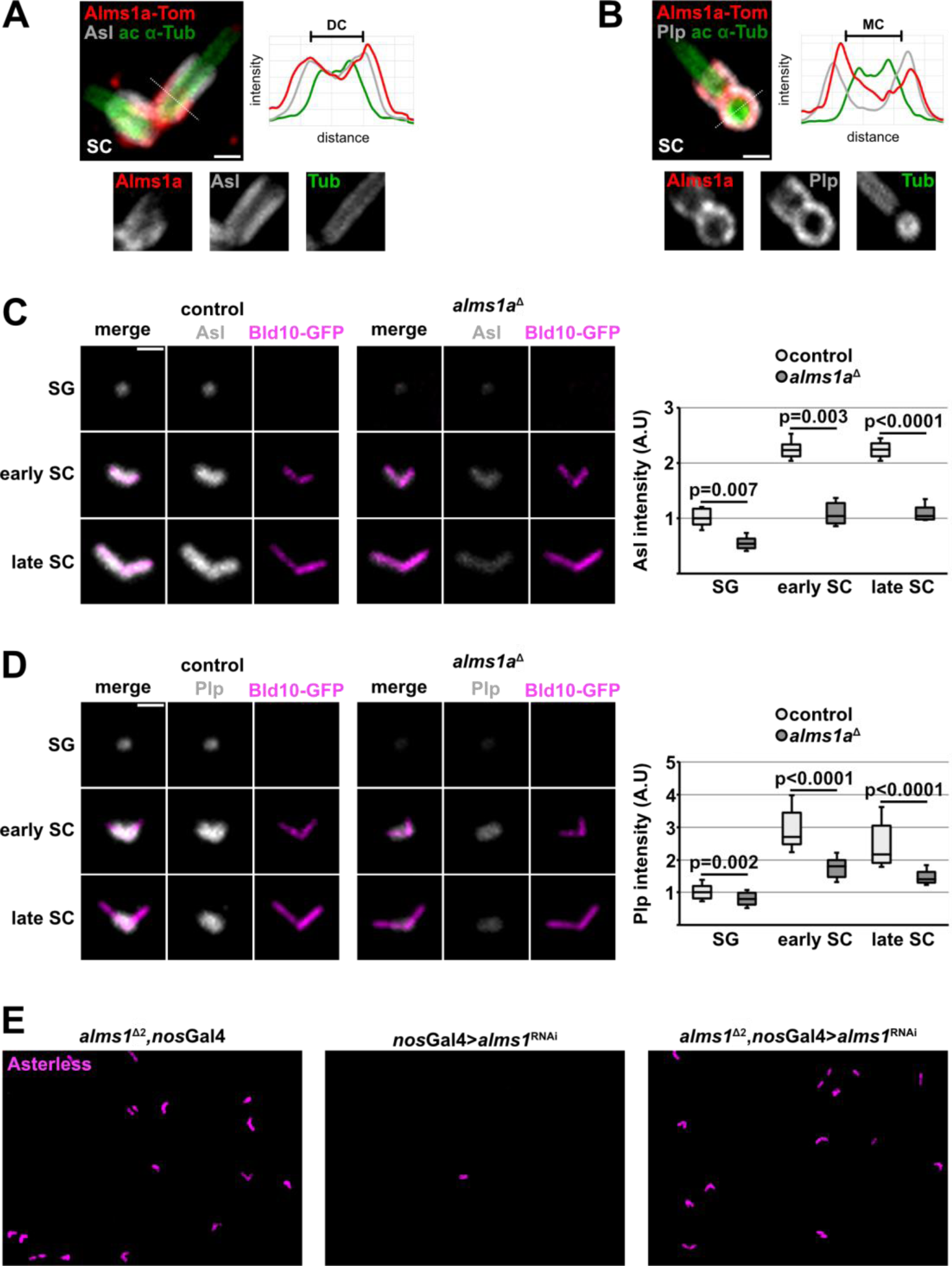
Alms1a is an inner PCM protein. U-ExM images of Alms1a-Tomato relative localisation with (A) Asl and (B) Plp. The fluorescence intensity along the dotted line is plotted. The mother (MC) or daughter centriole (DC) is indicated by a bracket. Scale bars, 1 μm. Standard images of (C) Asl and (D) Plp in w^1118^, Bld10-GFP (control) or *alms1*a^Δ^, Bld10-GFP and related fluorescence intensity quantification. Scale bars, 1 μm (E) Standard images of *nos*Gal4>*alms1*^RNAi^ in w^1118^ (middle) or *alms1*^2^^Δ^(right). Scale bars, 5 μm

We assessed the recruitment of Alms1 proteins in neuroblasts (NBs) of the larval brain. NBs are somatic stem-cells that generate the central nervous system of flies through asymmetric divisions ^23^. In NBs, Alms1a-Tomato localises at both mother and daughter centrioles, labelled with the PCM protein Asterless (Asl, Fig. 1D), with a higher concentration of Alms1a-Tomato on the mother compared to the daughter centriole in early interphase (Fig. 1D left, MC on the top, DC at the bottom). This asymmetry is still observed in pre-mitosis, after migration to the opposite pole of the NB of the mother centriole, which will be inherited by the differentiating cell (ganglion mother cell, GMC) ^23^ while the daughter centriole will be maintained in the NB (Fig. 1D right, DC #1, MC #2). In contrast to Alms1a, we were unable to detect Alms1b-GFP in the larval brain neither in NBs nor GMCs (not shown), in agreement with RNA-seq data showing low expression of *alms1b* in neurons and glial cells ^24^.

In syncytial embryos, which undergo 13 symmetric and synchronous mitosis ^25^, Alms1a-Tomato is observed at both centrioles from the first mitosis onwards. As in the male germline, we detected a faint Alms1a-Tomato staining at the onset of procentriole assembly in prophase (Fig. 1E, white, arrow). The signal becomes stronger on daughter centrioles at the beginning of the centriole-to-centrosome conversion in metaphase (Fig. 1E). In comparison, Alms1b-GFP was not detected in syncytial embryos and only observed at centrioles after cellularisation (not shown).

Thus, despite their high level of identity (more than 80%), Alms1a and Alms1b present very different spatiotemporal dynamics of centriolar recruitment in all tissues observed. These differences in centriolar recruitment and dynamics suggest distinct roles at the centriole. While Alms1b is only detected on mature centrioles (spermatocytes) or in cells which are not cycling or with a low cycling rate (post syncytial embryonic stages, absent in NBs), Alms1a is observed on all centrioles from the onset of procentriole assembly.

### Alms1a is an inner PCM protein

Alms1a-Tomato surrounds the centriolar wall, in a pattern reminiscent of the inner PCM proteins which are proposed to link centrioles and PCM and are hence also called bridge PCM proteins ^26,27^. Indeed, spatial analysis of staining intensity in U-ExM shows that Alms1a-Tomato colocalises with the two inner PCM proteins Cep152/Asterless (Asl) and PCNT/Pericentrin-like protein (Plp) (Fig. 2A,B). While Alms1a-Tomato fully surrounds the minute centrioles of spermatogonia (Fig. 1B, SG), it remains restricted to the proximal third of the growing centriole of spermatocytes as observed for Plp localisation (Fig. 1B, SC; Fig. 2B). Furthermore, Alms1a-Tomato localisation as a sleeve around the centriole is lost following cold treatment of the testis (data not shown) as previously observed for the PCM proteins. Altogether, these observations suggest that Alms1a is a novel inner PCM protein.

To position Alms1a in the hierarchy of PCM recruitment, we generated fly lines carrying deletion of *alms1a* by CRISPR/Cas9 (*alms1a*^Δ^, Fig. S1) and evaluated the consequences of *alms1a* loss of function on PCM protein. We quantified the fluorescence intensity of Asl and Plp at centrioles in control and *alms1a*^Δ^ spermatogonia, early and late spermatocytes. We observed a significant decrease of both Asl and Plp fluorescence intensity in *alms1a*^Δ^ cells, from all stages (Fig. 2C,D; Asl : control n=7 testes, *alms1a*^Δ^ n=8 testes; d-Plp : control n=8 testes, *alms1a*^Δ^ n=8 testes). Alms1a is thus required to either efficiently recruit or stabilise the inner PCM proteins Asl and Plp at centrioles.

### Compensatory mechanisms are triggered upon chronic *alms1a, alms1b* loss of function

A previous study demonstrated that depletion of both *alms1a* and *alms1b* by RNAi in asymmetrically dividing GSCs results in major loss of centrosomes ^19^. Here, despite the full deletion of *alms1a* locus, we failed to detect any centriole duplication loss (not shown). To exclude a possible compensation of *alms1a* loss by *alms1b*, we generated flies carrying a genomic deletion of both *alms1a,b* genes (*alms1*^2^^Δ^, Fig. S1). Although a small proportion of centriole pairs showed premature disjunction of the centrioles (not shown), centriole duplication in these flies is largely normal (Fig. 2E left). In contrast, we confirmed that *alms1a,b* depletion by RNAi in the whole germline, using a *nanos*-Gal4 driver expressed in the GSC, results in the complete loss of centriole duplication ^19^ (Fig. 2E middle, only one or two single centrioles observed per testis). The discrepancy between the two extreme phenotypes resulting from the acute *alms1* depletion by RNAi or the chronic *alms1* loss in *alms1*^2^^Δ^ flies could be due to either an off-target effect of the *alms1*^RNAi^ used or to compensation mechanisms triggered by the chronic loss of *alms1a* and *alms1b*. To discriminate between these two possibilities, we performed *alms1* depletion by RNAi in *alms1*^2^^Δ^ mutants (Fig. 2E right). For an off-target effect, *alms1*^RNAi^ should induce identical loss of centrioles in WT and *alms1*^2^^Δ^ backgrounds. However, *alms1*^2^^Δ^ flies with or without induction of *alms1*^RNAi^ present the same phenotype, *i.e*. no centriolar loss. *alms1*^2^^Δ^ flies have thus developed compensatory mechanisms which alleviate the loss of Alms1a,b proteins, indicating that Alms1 proteins play a critical role in centriole biogenesis in the male spermatogenesis that can be highly buffered during development.

### Alms1 proteins are general regulators of centriole duplication

To determine whether this function in centriole duplication is a general feature of Alms1 proteins, we extended our study to later stages of centriole duplication in the male germline and to the two other tissues studied above (Fig. 3). We depleted both *alms1a* and *b* in these tissues by expressing the *alms1*^RNAi^ with the appropriate tissue specific Gal4 drivers. We used *bam*-Gal4 (*bag of marbles*) ^28^ to induce *alms1*^RNAi^ expression in symmetrically dividing spermatogonial cysts, between 4 and 16-cells stages ^22^ (Fig. 3A,B). While control spermatocytes (*bam*-Gal4>*lacZ*^RNAi^) contain two centrosomes with one centriole pair each, for a total of 4 centrioles per cell (4C, 100% of cells, n=36 cells, 3 testes), *bam*-Gal4>*alms1*^RNAi^ cells contain only one centriole (1C, 92,1% of cells, n=89 cells, 5 testes) or no centriole (0C, 6,7 % of cells; Fig. 3A). This distribution is consistent with a centriole duplication failure event in: i) all cells within 8-cells spermatogonial cysts (each giving 2 cells with one centriole after the 4^th^ round of mitosis); and ii) in some cells at the 4-cells stage (giving two cells without centrioles and two with one centriole after the last round of mitosis, Fig. S2B). More, using U-ExM, we confirmed that the remaining mother centrioles in *bam*-Gal4>*alms1*^RNAi^ fail to assemble procentrioles as visualised with acetylated α-tubulin (94,1% of MC, n=34 centrioles, 2 testes) and do not recruit PCM protein Asl at the duplication site (Fig. 3B). The remaining 5,9% are duplicated centrioles observed at the spermatogonia stage, probably in cells where the *alms1*^RNAi^ was not activated yet. Together, these findings demonstrate that Alms1 proteins are crucial for centriole duplication in *Drosophila* male germline and that this role is not restricted to GSCs as previously proposed ^19^ but applies to all dividing germline cells. Strikingly, we observe that all the remaining centrioles present a normal distribution of Alms1a (Fig. 3C), indicating that centriolar duplication events do not depend on the pool of Alms1 surrounding the mother centriole, but more likely rely on a new pool of Alms1a recruited before Sas-6 cartwheel formation (Fig. 1C). In agreement of these observations, the remaining centrioles also show a normal ultrastructure by EM (Fig. S5), while no procentriolar structures could be detected. Together, these observations indicate that Alms1a,b are involved in the initiation of each centriole duplication event in the male germline.

**Figure 3.**
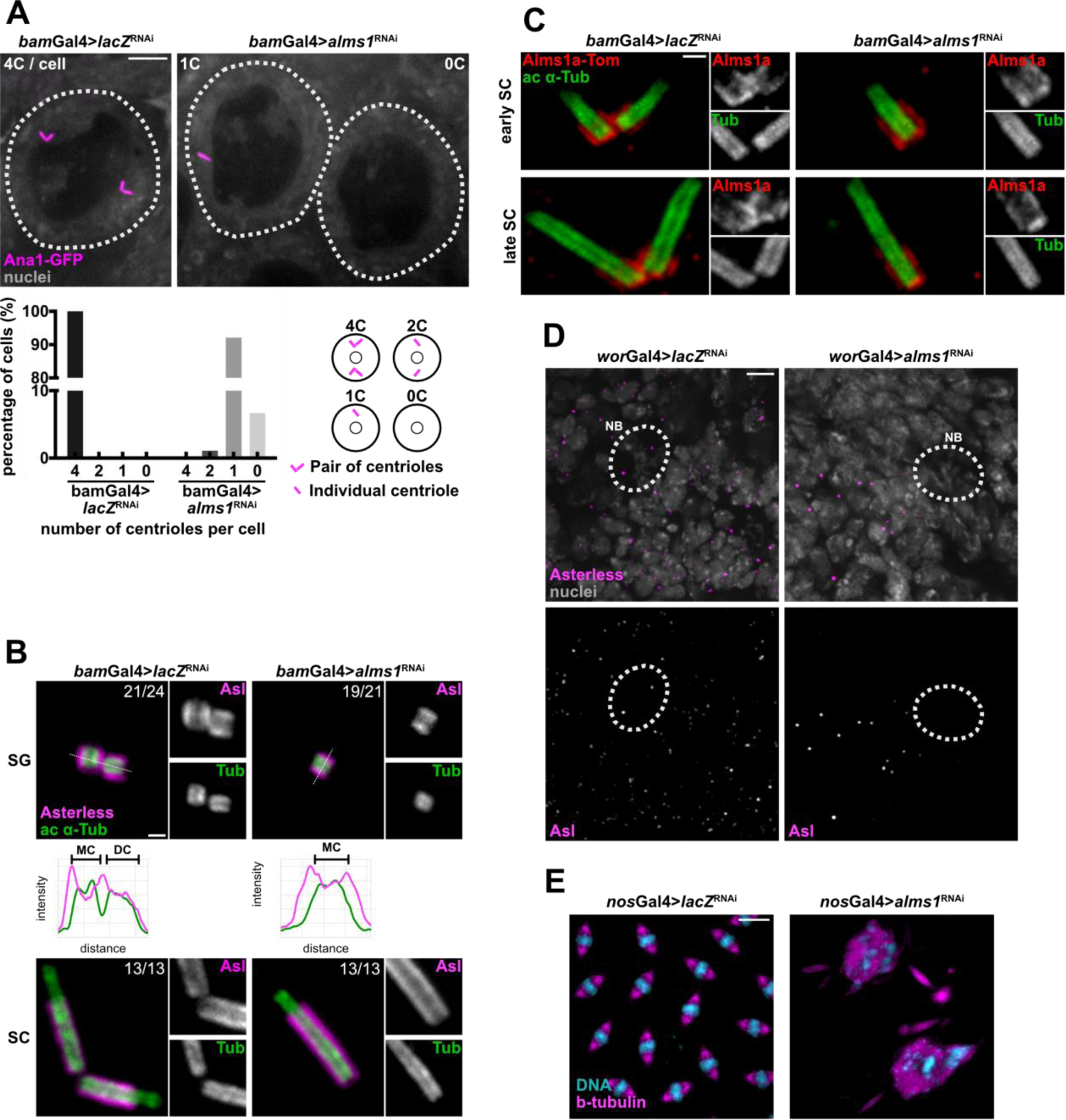
The acute depletion of Alms1 proteins lead to centriole duplication failure in various cell lineages. (A) Images of unexpanded (or conventionally fixed) *bam*-Gal4>*lacZ*^RNAi^ or *alms1*^RNAi^ spermatocytes (outline with white dotted lines) expressing Ana1-GFP (magenta) as a centriolar marker. Nuclei (gray). Quantification of the number of centrioles per spermatocytes (4C, 2C, 1C or 0C) in the indicated genotypes (*bam*-Gal4>*lacZ*^RNAi^ : n=36 cells, 3 testes; *bam*-Gal4>*alms1*^RNAi^ : n=89 cells, 5 testes). Scheme representing the observed distributions. Scale bars, 5 μm (B) U-ExM images of *bam*-Gal4>*lacZ*^RNAi^ or *bam*-Gal4>*alms1*^RNAi^ centrioles through spermatogenesis (spermatogonia SG; spermatocytes SC). Centriolar walls are revealed with acetylated α-tubulin antibody (green, ac a-tub). Asl (magenta) as a marker of PCM. On each image, numbers indicate the occurrence of the phenotype (*bam*-Gal4>*lacZ*^RNAi^ : n=37 centrioles, 2 testes; *bam*-Gal4>*alms1*^RNAi^ : n=34 centrioles, 2 testes). The fluorescence intensity along the dotted line is plotted. Mother centrioles (MC, on the left of all images) and, if necessary, daughter centrioles (DC, on the right) are indicated by brackets. Scale bars, 1 μm. (C) U-ExM images of Alms1a-Tomato (red) in *bam*-Gal4>*lacZ*^RNAi^ or *bam*-Gal4>*alms1*^RNAi^ centrioles. Scale bars, 1 μm. (D) Standard images of *wor*-Gal4>*lacZ*^RNAi^ or *wor*-Gal4>*alms1*^RNAi^ larval brain. NBs (outlined with white dotted lines) are identified based on the Miranda-GFP expression (not shown). Asl as a centriolar marker (magenta) and nuclei (gray). (E) Standard images of *nos*-Gal4>*lacZ*^RNAi^ or *nos*-Gal4>*alms1*^RNAi^ early embryos. β-tubulin for the mitotic spindle (magenta) and DNA (cyan). Scale bars, 5 μm.

To explore Alms1a,b function in the larval brain, we used the *worniu*-Gal4 driver (*wor*-Gal4) ^29^ to express the *alms1*^RNAi^. Control larval brains (*wor*-Gal4>*lacZ*^RNAi^) display many centrosomes in GMCs and 2 centrosomes in NBs (n=3 brains) while *wor*-Gal4>*alms1*^RNAi^ brains show a strong reduction of centrosomes in GMCs and a total loss of centrosome in NBs (n=5 brains, Fig. 3D), consistent with a role of Alms1 proteins in centriole duplication during NBs asymmetric division. We also generated embryos depleted in maternally provided *alms1* transcripts (from females expressing *alms1*^RNAi^ under the control of *nanos*-Gal4) ^30^, to investigate the contribution of Alms1a in symmetric division in somatic tissues. While control syncytial embryos (from *nanos*-Gal4>*lacZ*^RNAi^ females) undergo symmetrical and synchronous nuclear division cycles (n=3), *alms1*^RNAi^-derived embryos exhibit strong mitosis defects including multipolar spindles characteristics of centriole duplication failure (n=8; Fig. 3E).

Collectively, our results thus demonstrate that Alms1 proteins are required for centriole duplication in both symmetrically and asymmetrically dividing cells of the soma or germline.

### Alms1 proteins are required for cartwheel formation

The formation of the procentriole is the result of a complex cascade of phosphorylations initiated by the recruitment, concentration and transactivation of Plk4 as a single dot on the side of the mother centriole ^9,31^. Activated Plk4 then phosphorylates Ana2 in two steps: first on serine 38 (localised in the ANST motif at Ana2 N-terminus), allowing the efficient loading of Ana2 and the reinforcement of Plk4 localisation and activation at the duplication site ^6,11,12^, then on the STAN motif, triggering the recruitment of Sas-6 by Ana2 ^11,32,33^ and the formation of the cartwheel onto which the procentriole assembles (Fig. 4A).

**Figure 4.**
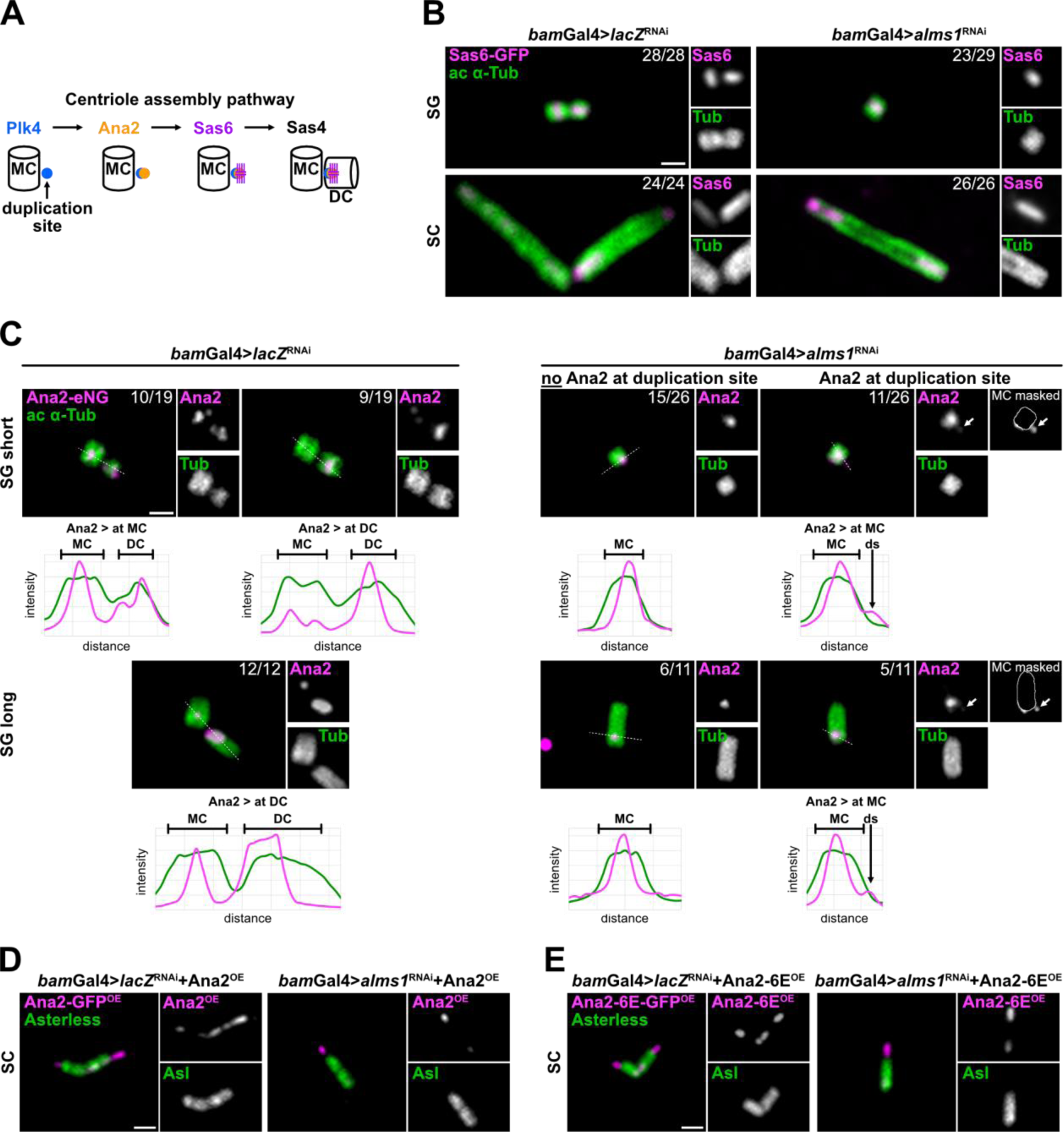
Alms1 proteins are essential for Sas-6 recruitment and required to stabilise Ana2. (A) Scheme representing part of the known protein hierarchy responsible for centriole duplication, leading to procentriole formation. (B) U-ExM images of Sas-6-GFP (in magenta) in *bam*-Gal4>*lacZ*^RNAi^ (control) or *bam*-Gal4>*alms1*^RNAi^ during spermatogenesis (spermatogonia SG; spermatocytes SC). Centriolar walls are revealed with acetylated α-tubulin antibody (green). On each image, numbers indicate the occurrence of the phenotype. (*lacZ*^RNAi^ : n=52 centrioles, 4 testes; *alms1*^RNAi^ : n=55 centrioles, 6 testes). (C) U-ExM images of Ana2-mNeonGreen knock-in (Ana2-eNG, in magenta) in *bam*-Gal4>*lacZ*^RNAi^ (control) or *bam*-Gal4>*alms1*^RNAi^. Centrioles at the spermatogonium stage are divided into two categories: SG short, when the length of the centrioles does not exceed their width and SG long, when their length exceeds their width. On each image, numbers indicate the occurrence of the phenotype (for SG and SC, *lacZ*^RNAi^: n=71 centrioles, 3 testes; *alms1*^RNAi^ : n=57 centrioles, 5 testes; SC on Fig. S3). The fluorescence intensity along the dotted line is plotted as a function of distance on the graphs. Mother centriole (MC, on the left of all images) and daughter centriole (DC, on the right) are indicated by an interval, duplication site (ds) by an arrow. Scale bars, 1 μm. (D-E) Standard images of *bam*-Gal4> *lacZ*^RNAi^ or *bam*-Gal4>*alms1*^RNAi^ spermatocytes overexpressing either (D) Ana2-GFP^OE^ (*lacZ*^RNAi^ : n=2 testis; *alms1*^RNAi^ : n=5 testis) or (E) constitutively active Ana2 (Ana2-6E-GFP^OE^) (*lacZ*^RNAi^ : n=2 testes; *alms1*^RNAi^ : n=5 testes). Scale bars, 1 μm.

Sas-6-GFP is detected at the proximal end of the mother centriole in both *alms1*^RNAi^ and control conditions (U-ExM, Fig. 4B), in agreement with the conservation of the cartwheel after centriole elongation in *Drosophila* ^2^. In control testes, a dot of Sas-6-GFP is also observed on the side of most mother centrioles initiating centriole duplication (n=12/15 centrioles, 4 testes, not shown) and subsequently in the proximal lumen of all growing procentrioles (Fig. 4B, *bam*-Gal4> *lacZ*^RNAi^, n=52 centrioles, 4 testes). In contrast, we never observed Sas-6-GFP affixed to mother centrioles in late spermatogonia nor in spermatocytes in *bam*-Gal4>*alms1*^RNAi^ (Fig. 4B, n=49 centrioles, 6 testes). The only occurrences of Sas-6-GFP affixed to mother centrioles were observed in young spermatogonia, in which the *bam*-Gal4 is likely not active yet (n=6 centrioles, not shown). Hence, in absence of Alms1, the cartwheel of Sas-6 fails to assemble at the duplication site.

### Alms1 proteins are required for Ana2 stabilisation at the centriole duplication site

We hypothesise that the lack of cartwheel in *alms1*^RNAi^ might reflect a role of Alms1 either in the recruitment of Ana2 at the site of centriole duplication or in the recruitment of Sas-6 by Ana2.

We thus characterised Ana2 localisation in control and *alms1*^RNAi^ conditions by U-ExM, using a Ana2-mNeonGreen knock-in line (Ana2-eNG) ^34^. In control spermatogonia, Ana2-eNG localises at the proximal inner end of the mother centriole and forms a dot at the base of the forming procentriole (Fig. 4C, left) ^6,35^. As daughter centrioles elongate, we observed a shift of Ana2 abundance from the mother to the daughter centriole: while Ana2 is more abundant at the mother centriole than at the nascent procentriole in the early duplicating centrioles (55% of SG short centrioles, 10/19 pairs), it becomes more abundant at daughter centrioles as they grow (SG long ; SC, Fig. S3) to finally fully disappear from some mother centrioles (n=8/32 SC, Fig. S3) ^11,35^.

In *alms1*^RNAi^ condition, Ana2-eNG is also nestled in the proximal end of all remaining mother centrioles in spermatogonia and spermatocytes. We failed to detect any Ana2-eNG on the side of 58% of the un-duplicated centrioles observed (Fig. 4C, middle, 15/26; Fig. S3), whereas for 42% we observed a very faint Ana2-eNG staining at the centriole duplication site, almost always less abundant than the concentration at the base of the “mother” centriole (Fig. 4C, right; Fig. S3). Altogether our observations thus suggest that Alms1 is involved in the efficient recruitment or stabilisation of Ana2 at the duplication site and that this reduced Ana2 activity is responsible for the lack of Sas-6 recruitment. To trigger cartwheel formation, Ana2 needs to be activated by phosphorylation of its STAN domain by Plk4 ^6,32,36,37^. We thus aimed to determine if the lack of centriole duplication in *alms1*^RNAi^ condition could be rescued by over-expression of Ana2-GFP either WT or with a phospho-mimetic STAN domain (respectively Ana2-GFP^OE^, Ana2-6E-GFP^OE^) ^38,39^. However, overexpressing Ana2-GFP or Ana2-6E-GFP in *alms1*^RNAi^ spermatogonia does not rescue the duplication defect (*bam*Gal4>*alms1*^RNAi^+Ana2-GFP^OE^, Fig. 4D; +Ana2-6E-GFP^OE^, Fig. 4E). These results lead us to propose that centriole duplication failure is directly related to reduced recruitment of Ana2 at the duplication site.

### Alms1 proteins are required for Plk4 activity

Ana2 recruitment at the procentriole is under the direct control of Plk4 ^6^. In wild-type conditions, PlK4 activity is tightly buffered to avoid the formation of supernumerary duplication sites and hence centriole overduplication. This control can be overcome by either Plk4 overexpression or expression of non-degradable Plk4 ^6,10,40^. Hence, in *Drosophila* spermatogonia, increase of Plk4 generates extra-centrioles organised in rosettes in the centrosome and *de novo* centriole formation ^9^. In agreement with this latter result, Plk4 overexpression in our control background (Plk4^OE^, *bam*-Gal4>*lacZ*^RNAi^) induces both rosettes containing more than three centrioles (3^+^C, Fig. 5A) and centrioles formed *de novo* (not shown). In contrast, we almost only observe un-duplicated centrioles when Plk4 is overexpressed in spermatogonia depleted in *alms1* (Plk4^OE^, *bam*-Gal4>*alms1*^RNAi^, Fig. 5A) and an extremely low percentage of centriole pairs (less than 1%, n=9 testis). However, endogenous Plk4 (Plk4-eGFP) ^41^ concentrates at the base of all procentrioles in control conditions (Fig. 5B*, bam*-Gal4>*lacZ*^RNAi^, n=33 centrioles, 3 testes) and forms a dot on the side of 91,7% of the un-duplicated centrioles in spermatogonia depleted in Alms1 (Fig. 5B, *bam*-Gal4>*alms1*^RNAi^, n=72 centrioles, 6 testes; SC in Fig. S4).

**Figure 5.**
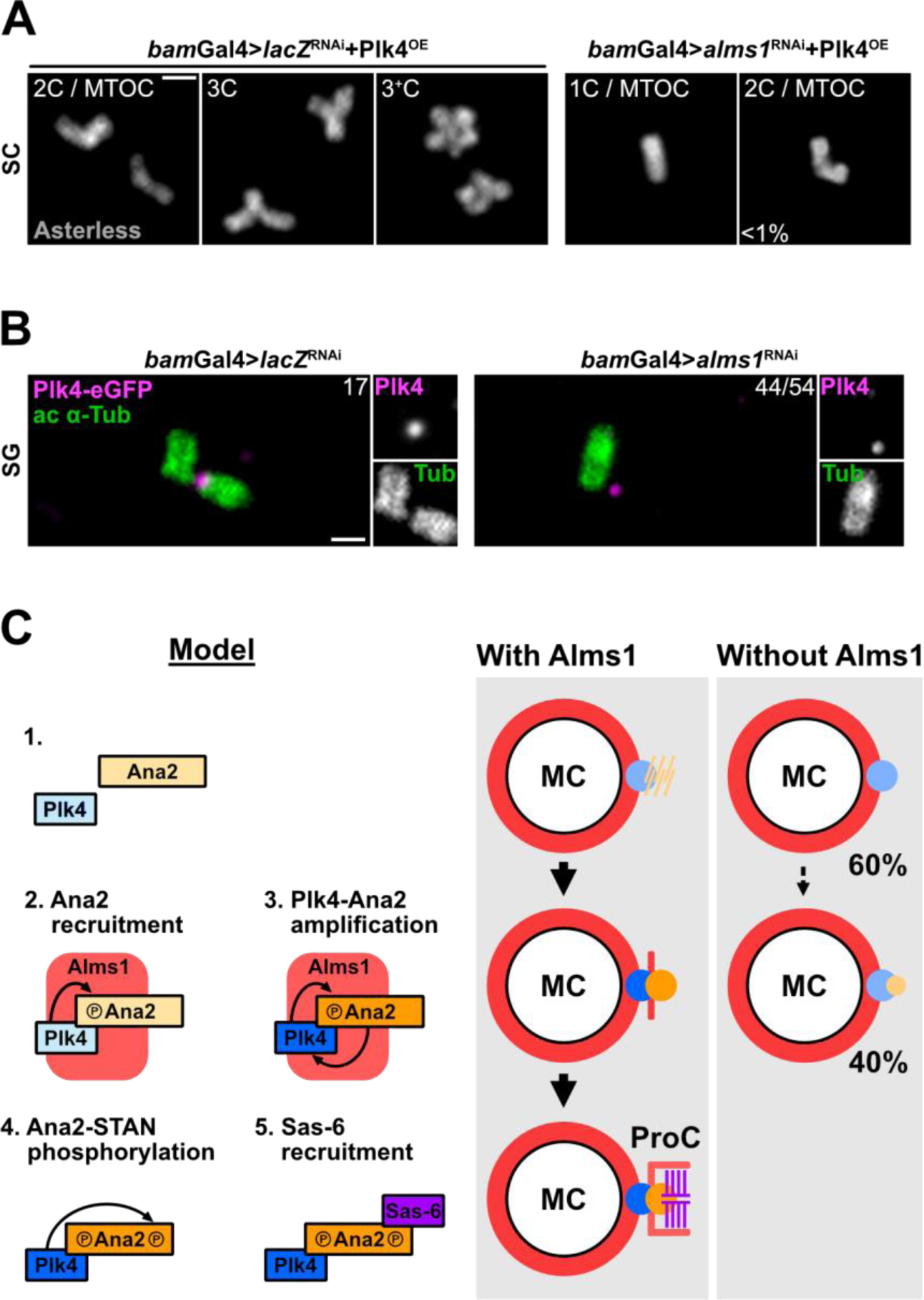
Alms1 proteins act downstream of Plk4. (A) Images of unexpanded *bam*-Gal4>*lacZ*^RNAi^ or *bam*-Gal4>*alms1*^RNAi^ spermatocytes (SC) overexpressing Plk4 (Plk4^OE^, not tagged). Asterless (grey) as a centriolar marker. The different phenotypes observed are shown by genotype: 1, 2, 3 centrioles or more (3+) per centrosome (MTOC). (*lacZ*^RNAi^ : n=7 testes; *alms1*^RNAi^ : n=9 testes). Scale bars, 1 μm. (B) U-ExM images of Plk4-meGFP knock-in (in magenta) in *bam*-Gal4>*lacZ*^RNAi^ (control) or *bam*-Gal4>*alms1*^RNAi^in spermatogonia (SG). Centriolar walls are revealed with acetylated α-tubulin antibody (green). On each image, numbers indicate the occurrence of the phenotype (For SG and SC, *lacZ*^RNAi^ : n=33 centrioles, 3 testes; *alms1*^RNAi^ : n=72 centrioles, 6 testes). Scale bars 1 μm. (C) Proposed model for Alms1 role during centriole duplication: Alms1 could contribute to the recruitment of Ana2 by Plk4 (step 2) and/or the subsequent positive amplification loop of Plk4-Ana2 concentration and activity (step 3). Alms1 could be contribute to bridging Plk4 and Ana2, thus creating a proximity allowing the positive feed-back loop. Lack of Alms1 would prevent Ana2 full (or efficient) recruitment by Plk4, thus resulting in centriole duplication failure.

These experiments demonstrate that Alms1a,b are not necessary for Plk4 recruitment on centrioles but could be required for Plk4 activity, placing Alms1 downstream of Plk4 but upstream of Ana2. Thus, we propose that Alms1 enhances Plk4-Ana2 functional interactions and that in absence of Alms1 the Plk4-Ana2 reinforcement loop is impaired, preventing the concentration of active Ana2 and leading to dramatic loss of Sas-6 assembly and of nascent procentriole formation (Fig. 6).

## Discussion

Here we show that, in *Drosophila*, Alms1 proteins are general regulators of centriole duplication as their acute depletion by RNAi results in complete failure of centriole duplication (Fig. 3), not only in asymmetrically dividing germline stem cell ^19^ but in all asymmetrically and symmetrically dividing somatic and germline cells analysed. More, we demonstrate that Alms1a,b are either essential (following acute RNAi depletion) or dispensable (upon chronic loss) intermediate players between PlK4 and Ana2, such apparent contradictory properties illustrating striking buffering capacities of cells or tissues to maintain centriole number during development.

Indeed, centriole duplication occurs only once per cell cycle to generate exactly two pairs of centrioles that will be inherited by the two daughter cells. This process is tightly regulated to ensure that duplication occurs at the right time and generates exactly two new pairs of centrioles. Indeed, alterations of centriole number or maturation lead to defects in cell cycle and are associated with numerous pathologies including cancers or microcephaly ^2^. Extensive screens in various models and in particular in *C. elegans* and *Drosophila* led to the identification of a few conserved proteins that are at the core of the centriole duplication process: the Plk4/Sak kinase (in *D.m*., PLK4 in mammals, ZYG-1 in *C.e*.), Ana2 (STIL, SAS-5), Sas-6 (SAS-6, SAS-6), Spd2 (CEP192, SPD-2) and Sas-4 (CPAP, SAS-4) ^32,39,42–45^, but failed to identify Alms1 proteins.

The main explanation why ALMS1 has remained unidentified so far is the striking difference between acute and chronic loss of Alms1a,b proteins that we have observed in flies. We do not have a molecular explanation of the compensatory mechanisms at work, but several observations in the literature support the hypothesis that centriole duplication is very sensitive to variations in the expression of core molecular players. In particular, it has been shown that overexpression of centriole duplication factors can induce centrosome amplification and centriole overduplication ^46,47^. Thus, we anticipate that during chronic loss of Alms1a and b, the expression of some core players and/or other components still to be identified are modulated to fine tune centriolar duplication. Future investigations of the compensatory mechanisms involved in *alms1* loss of function will likely provide fascinating knowledge on centriole homeostasis in cells. Such compensatory mechanisms have already been described in other cellular processes in whole organisms ^48^ and thus represent a challenging complexity for the understanding of these processes. In all our chronic loss experiments, we could not discriminate between the function of Alms1a and Alms1b. Indeed, due to the extremely strong conservation of their transcript sequence, we could not successfully design specific shRNAi. However, based on their dynamics of expression, Alms1a being recruited just prior to cartwheel formation whereas Alms1b is recruited after the initiation of procentriole formation, we anticipate that Alms1a is the one involved in the duplication process. More specific genetic strategies will however need to be designed to formerly determine the respective contribution of Alms1a or Alms1b in controlling centriole duplication.

We demonstrate that Alms1a,b proteins play a crucial role in centriole duplication and we show that Alms1a,b are novel regulators of cartwheel formation by modulating Ana2 stabilisation (Fig. 4, Fig.6). Ana2 stabilisation at the centriolar duplication site is a complex process that relies on successive phosphorylation of Ana2 by Plk4 and at least two reinforcement loops to stabilise both Plk4 and Ana2 and initiate Sas-6 recruitment in a single dot. It has been shown in various cell culture systems that centriole duplication starts with the recruitment of Plk4, as a ring surrounding the mother centriole ^6,49^. The binding of Ana2/STIL ^6,49^ protects Plk4 from auto-phosphorylating its degron motif which triggers Plk4 degradation by the ubiquitin-proteasome pathway ^50–52^. As a consequence, Plk4 is degraded in the whole ring with the exception of the single Plk4/Ana2 interaction site. There, a positive reinforcement loop is initiated whereby Plk4 phosphorylation of Ana2 stabilises it and in turn phosphorylated Ana-2 stabilises activated Plk4 ^6,12,35,37^. This loop results in phosphorylation of Ana2 on its STAN domain, making it competent to recruit Sas-6 and trigger the formation of the cartwheel. In contrast to isolated mammalian or *Drosophila* cells, we never observed Plk4 forming a ring around the centriole in *Drosophila* tissues but we directly observed a single dot of Plk4 at the base of the procentriole from the very early steps of centriole duplication (Fig. 5B). In these tissues, Ana2, as observed with U-ExM, forms a more diffuse dot (Fig. 4C) than Sas-6 (Fig. 4B). Following acute depletion of Alms1, we still observe Plk4 at the duplication site of all centrioles (Fig. 5B) but we fail to detect Ana2 in 58% of the centrioles and it is reduced in the remaining 42% (Fig. 4C). Centriole duplication failure after *alms1*^RNAi^ depletion is not rescued by the expression of phosphomimetic Ana2, nor by increasing the overall amount of Ana2, suggesting that other limiting steps than Ana2 activation on its STAN motif are responsible for defective Ana2 stabilisation in Alms1 depleted cells. Stabilisation of Plk4/Ana2 has also been proposed to rely on Sas-4/CPAP which serves as a platform bringing the two proteins in proximity ^6,12,35^. Interestingly, Alms1 has been identified in BioID screens as a potential interactor of CPAP ^53,54^ and is a direct interactor of Plk4 ^19^. One can thus speculate that, as proposed for Sas-4/CPAP, Alms1 could be involved in stabilising Ana2 at the duplication site by bridging together Plk4 and Ana2 (Fig. 6). Future experiments will be required to test this hypothesis and position Alms1 with respect to the Sas-4/PLK4/Ana2 module. Together, our work indicates that Alms1a,b in flies are critical regulators of centriole duplication which function can be compensated.

If we do not observe centriolar defects after chronic loss of Alms1a,b, we however do observe centriole disjunction, a phenotype also observed in human cells after RNAi depletion of ALMS1 ^16^, thus indicating that this function of Alms1 proteins is likely conserved across evolution. On the other hand, acute loss of ALMS1 is not sufficient to reveal centriolar duplication defects in mammalian cells. It is thus difficult to evaluate to which extent the function of ALMS1 in centriole duplication is conserved in mammals. Whereas there are only two ALMS1 domain containing proteins in flies, there are 3 proteins containing an ALMS domain in humans, ALMS1, FATS and CEP295 ^16^. No centriolar functions have been described for FATS to date but experimental data indicate a role for CEP295 in centriole elongation ^55^. The three proteins thus seem to have acquired different specialised functions, but experimental evidences are lacking to evaluate possible redundant roles in the ALMS domain family of proteins.

In conclusion, our work shows that Alms1 proteins are general regulators of centriole duplication in flies, acting during the complex, Plk4-dependent, stabilisation step of Ana2 to initiate cartwheel formation. It also reveals striking, but still to be understood, buffering capacities of cells and tissues during development to compensate for *alms1* loss of function. Understanding if Alms1 function in centriole duplication is conserved in humans is a future challenge as *Alms1* is the only gene associated with the extremely rare Alström syndrome in humans ^17,18^.

## Methods

### Fly stock and husbandry

Flies were raised at 18°C, 25°C or 29°C on a standard nutrient medium (cornmeal, yeast, agar, nipagin, ethanol). Fly lines generated and stocks used are listed in Table 1.

**Table 1.** Drosophila melanogaster stocks.

|  |  |  |
| --- | --- | --- |
| **Alms1a-Tomato**<br>**in rescue condition** | *alms1a<sup>Δ</sup>*/FM7h; ; Alms1a-Tomato/TM3, Ser | This study |
| ***alms1a<sup>Δ</sup>*** | *alms1a<sup>Δ</sup>*/FM7h; ; | This study |
| **Alms1a-Tomato** | ; ; Alms1a-Tomato/TM3, Ser | This study |
| **Alms1b-GFP**<br>**in rescue condition** | *alms1b<sup>Δ</sup>*/FM7; Alms1b-GFP/CyO; | This study |
| ***alms1b<sup>Δ</sup>*** | *alms1b<sup>Δ</sup>*/FM7; ; | This study |
| **Alms1b-GFP** | ; Alms1b-GFP/CyO; | This study |
| ***alms1<sup>2Δ</sup>*** | *alms1a<sup>Δ</sup>,alms1b<sup>Δ</sup>*/FM7h; ; | This study |
| ***lacZ<sup>RNAi</sup>*** | *w<sup>1118</sup>*; P{GD936}v51446; | VDRC 51446 |
| ***alms1<sup>RNAi</sup>*** | ; UAS-P{TriP.HMJ30289}/CyO; | BL 63721 |
| ***bam*-Gal4** | ; If/CyO; *bam*-Gal4, UAS-Dcr2/TM6B |  |
| ***bam*-Gal4** | *bam*-Gal4, UAS-Dcr2/FM7h; ; |  |
| ***nos*-Gal4** | *w<sup>1118</sup>*; ; P{w[+mC]=GAL4::VP16-<br>*nanos*.UTR}CG6325[MVD1] | BL 4937 |
| ***wor*-Gal4, Miranda-GFP** | *w<sup>\*</sup>*; P{w[+mC]=*wor*.GAL4.A}2, P{w[+mC]=UAS-<br>*mira*.GFP}1.2/CyO ; Dr/TM6B | Modified from BL 56555 |
| **Ana1-GFP** | *w<sup>\*</sup>*; Bl/CyO ; P{ana1-GFP.B}/TM6B |  |
| **Bld10-GFP** | ; UASp-endo-Bld10-GFP [142.1]/CyO; | from T. Megraw <sup>61</sup> |
| **Plk4-eGFP** | *w<sup>\*</sup>*; If/CyO; meGFP-Plk4/TM6B | from M. Bettencourt-Dias <sup>41</sup> |
| **Ana2-eNG** | *w<sup>\*</sup>*; eAna2-mNG 3.72/SM5; | from J. Raff <sup>34</sup> |
| **Sas-6-GFP** | *w<sup>\*</sup>*; pUbq-GFP-Sas-6/CyO; | from R. Basto |
| **Plk4<sup>OE</sup>** | ; UASp-Plk4/CyO; | From M. Bettencourt-Dias |
| **Ana2-GFP<sup>OE</sup>** | ; pUbq-GFP-Ana2/SM5; | from J. Raff <sup>62</sup> |
| **Ana2-6A-GFP<sup>OE</sup>** | ; pUbq-Ana2-6A-GFP/SM6a; | from J. Raff <sup>38</sup> |
| **Ana2-6E-GFP<sup>OE</sup>** | ; pUbq-Ana2-6E-GFP/SM6a; | from J. Raff <sup>38</sup> |

### Plasmids and *Drosophila* gene constructs

All primer sequences are available upon request. All transgenic constructs were injected by BestGene Inc.

### *Drosophila* Alms1 reporter gene constructs

Alms1a-Tomato was obtained by cloning the PCR product (F-Alms1a-Tom/R-Alms1a-Tom), containing 1.73 kb upstream regulatory sequences and the entire coding sequence (4,23 kb), in frame with Tomato in BglII-NotI sites of pJT108 ^56^.

For Alms1b-GFP construct, the PCR product (F-Alms1b-GFP/R-Alms1b-GFP) including 1,5 kb upstream regulatory sequences and the entire coding sequence (3,3 kb) was cloned in frame with GFP in the BglII site of pJT61 ^56^ using Gibson Assembly Master Mix (New England Biolabs Inc.)

Alms1a-Tomato was integrated in the 89E11 VK00027 landing site on the third chromosome and Alms1b-GFP on the 53B2 VK00018 landing site on the second chromosome by PhiC31 integrase (BestGene Inc.).

### Generation of *alms1a*^Δ^, *alms1b*^Δ^ and *alms1*^2^^Δ^ by CRISPR/Cas9

*alms1a*^Δ^ allele was generated by CRISPR/Cas9 induced deletion (NHJE). Three couples of gRNAs (gRNA1Alms1a+gRNA2Alms1a, gRNA3Alms1a+gRNA4Alms1a and gRNA5Alms1a+gRNA6Alsm1a) were respectively cloned in pBFv-U6.2B vector ^57^ and injected together in CG12179[NP]/(FM7h);;vasa-Cas9 embryos. Flies were crossed to Hira[-]/FM7h females or FM7h/Y males and offspring were selected for white-eyed-flies. Deletion in the *alms1a* locus was further characterised by PCR with primers F-Alms1aKO/R-Alms1aKO and confirmed by sequencing.

*alms1b*^Δ^ allele was generated by CRISPR/Cas9 induced homologous direct repair ^58^. The 1.5 kb 5’ homology arm and 1.4 kb 3’ homology arm were amplified by PCR (F-5’armAlms1b/R-5’armAlms1b and F-3’armAlms1b/R-3’armAlms1b) on genomic DNA from vasa-Cas9 flies and cloned, respectively, into the NheI-KpnI sites and BglII-AvrII sites of pJT38 plasmid (pRK2 plasmid ^59^ containing an attB cassette). Two gRNAs (gRNA1Alms1b and gRNA2Alms1b) were cloned in the pBFv-U6.2B vector ^57^. The two constructs were injected in vasa-Cas9 embryos. Flies were crossed to Hira[-]/FM7h females or FM7h/Y males and the offspring were screened for red-eyed-flies. Homologous recombination was checked by PCR (F-5’Alms1bKO/R-5’Alms1bKO and F1-3’Alms1bKO/R-3’Alsm1bKO).

*alms1*^2^^Δ^ allele was generated from *alms1a*^Δ^ flies by CRISPR/Cas9 induced homologous direct repair on the *alms1b* locus. Template repair vector was constructed as above, with the exception of the 5’ arm, which was obtained by PCR with primers F-5’armAlms1a^Δ^/R-5’armAlms1b on genomic DNA from *alms1a*^Δ^ flies. This new template repair was injected together with the previous pBFv-U6.EB vector expressing gRNA1Alms1b and gRNA2Alms1b in *alms1a*^Δ^ embryo. Flies were crossed to Hira[–]/FM7h females or FM7h/Y males and the offspring were screened for red-eyed-flies. Homologous recombination was checked by PCR (F2-5’Alms1bKO/R-5’Alms1bKO and F2-3’Alms1bKO/R-3’Alms1bKO).

### Classical immunofluorescence preparations

#### Testes

Testes from young adults were dissected in PBS and fixed 20 min in PFA 4%. For each fly, a single testis was kept to avoid analysing the same individual twice. After 20 min of permeabilisation in PBS-Triton X-100 0.1% (PBST), testes were blocked 1h in PBST/BSA 2% (PBST-BSA) and then incubated with primary antibodies (in PBST-BSA, see Table 2) overnight at 4°C. After 4 washes, testes were incubated 2h with appropriate secondary antibodies and Hoechst. After 4 washes, testes were mounted in Vectashield Antifade Mounting Medium. If immunofluorescence was not required, testes were incubated 2h in Hoechst in PBST.

**Table 2.** Primary antibodies used.

|  |  |  | Classical preparation | U-ExM |
| --- | --- | --- | --- | --- |
| **Anti acetylated α-Tubulin** | mouse, IgG2b | clone 6-11B-1, T6793, Sigma-Aldrich, lot 017M4806V | - | 1:500 |
| **Anti RFP** | mouse, IgG2c | 6G6, ChromoTek, 6G6, lot 107272-07-02 | - | 1:100 |
| **Anti GFP** | rabbit | TP401, Chemokine, lot 081211 | - | 1:100 |
| **Anti GFP** | rabbit | 632459, Living Colors, lot1105001 | 1:500 | - |
| **Anti mNeonGreen** | rabbit | Cell Signaling Technology | - | 1:100 |
| **Anti Asterless** | rat | from G. Rogers | 1:75000 | 1:5000 |
| **Anti Plp** | rabbit | from G. Rogers | 1:2000 | 1:1000 |
| **Anti β-tubulin** | mouse, IgG1 | E7, Developmental Studies Hybridoma Bank | 1:100 | 1:500 |

#### Brains

Larva collection was performed as follows: 20% sucrose solution was added in tubes containing larvae in food. Supernatant containing larvae was collected, rinsed in PBS and larvae were placed on a petri dish with agar. Third instar larvae were selected for dissection. Brains were dissected in PBS during sessions not exceeding 20 min and then fixed 25 min in PFA. After 2 baths of 10 min each in PBST 0.3%, brains were blocked 2h in PBST/BSA 2% and then incubated with anti α-tubulin overnight at 4°C. After 4 washes, brains were incubated with the appropriate secondary antibody and Hoechst overnight at 4°C. After 4 washes, brains were mounted in Vectashield Antifade Mounting Medium.

#### Embryos

30min-2h30 synchronised embryos were collected on a petri dish with agar. Embryos were dechorionated for 5 min in bleach, devitelinised by vigorous shaking in heptane-methanol solution (1:1 ratio) and fixed in methanol. Embryos were blocked in two 30 min baths of PBST 0.1%/BSA 3% and incubated in anti β-tubulin overnight at 4°C. After 4 washes, embryos were incubated with the appropriate secondary antibody and Hoechst overnight at 4°C. After 4 washes, embryos were mounted in Vectashield Antifade Mounting Medium.

### PCM proteins fluorescence quantification

w^1118^ and *alms1a*^Δ^ testes were prepared for a classical immunofluorescence and incubated with anti-Asterless or anti-Plp antibodies. Samples of the different genotypes were processed identically and simultaneously. Adjustments of acquisition parameters were performed on one testis isolated from each genotype. Acquisitions were then made with the same exposure and laser parameters for all testes. Fluorescence quantification was performed after sample anonymisation by a third party. For each testis, Asl or Plp and Bld10-GFP fluorescence was quantified for 10 centrioles per stage (spermatogonia, early spermatocytes, late spermatocytes) using a threshold to delimit the area to be quantified and from which background noise was subtracted (macros available upon request). Two independent quantifications were performed for both Asl and Plp. For Asl, quantification 1 : control n=5 testes, *alms1a*^Δ^ n=5 testes ; quantification 2 : control n=2 testes, *alms1a*^Δ^ n=3 testes. For Plp, quantification 1 : control n=5 testes, *alms1a*^Δ^ n=5 testes ; quantification 2 : control n=3 testes, *alms1a*^Δ^ n=3 testes.

**U-ExM preparations** (adapted from ^20^).

### Testes

#### Crosslinking

Testes from young adults were dissected in PBS and incubated PBS/1.4% Formaldehyde - 0.3% Acrylamide (PBS-FA-AA) overnight at 18°C. Prior to the gelation, testes were incubated 2h in the monomer solution (PBS/19% Sodium Acrylate - 10% Acrylamide - 0.1% Bis-Acrylamide - 0.5% TEMED – Hoechst 1/500) to allow reagents to penetrate the inner layers of the tissue. Testes were then deposited in packs of 3-4 on 12mm coated coverslips (0.2 mg/ml Poly-D-Lysine during 2h at 37°C) and all the monomer solution was absorbed with filter paper. Coverslips were deposited on a silicone cushion made in 50 ml Falcon tube and spun down using a swing rotor for 5 min at 5000 rpm at 10°C. 38μL of cold monomer solution with 0.5% APS was deposited on coverslips which are immediately put upside down on a parafilm tensed on a microscope slide laid on an ice-cold block. Gelation was started on cold for 5 min before transfer for 1h at 37°C.

#### Relaxation of mechanical strains

Gels were cut around groups of testes on the coverslip using a biopsy punch (4mm diameter). Punches were incubated in denaturation buffer (SDS 200mM – NaCl 200mM – Tris pH9 50mM) for 1.5 h at 95°C. After 3 washes in water, gels were preserved in PBS at 4°C.

#### Immunofluorescent labelling

Gels were blocked 2h in PBS/Tween 0.05% - BSA 1% - Sodium azide 0.02% (PBSTw-BSA) and then incubated with primary antibodies (in PBSTw-BSA) in a humid chamber overnight at 4°C. After 4 washes, gels were incubated overnight with appropriate secondary antibodies and Hoechst and washed again.

#### Expansion

Gels were expanded for 2h in 2 baths of water. Gels were mounted on 15mm coated coverslips (0.2 mg/mL PolyD-Lysine during 2h at 37°C).

### Brains

Larvae were collected and selected as described above. Brains were dissected in PBS and immediately incubated in PBS-FA-AA. PBS-FA-AA solution was renewed before incubation overnight at 18°C. Brains were then dissected again on 12mm coated coverslips to remove the two optic lobes. The rest of the protocol is identical as described above.

#### Embryos

Embryos were collected, dechorionated and devitenilised as described above. The rest of the protocol is identical to the testes one.

### U-ExM controls

Expansion factor is calculated in each experiment by measuring the size of the gel disk obtained by standardised biopsy punch (4mm diameter) after expansion. Centrioles expansion isotropy and factor was validated by measuring centriole diameter (electron microscopy reference: 200nm ± 12nm).

### Image acquisition

All images (classical and U-ExM) were obtained using IX 83 inverted microscope from Olympus, equipped with a Yokagawa CSU-X1 Spinning Disk Unit, Borealis technology for homogeneous illumination and Ixon3 888 EM-CCD camera from Andor. The oil immersion Plan Apochromat 60x / 1.42 NA objective from Olympus was used for all acquisitions. All images were processed and analysed with FiJi ^60^.

### Transmission Electron Microscopy

Testes from pupae and 3-4 day old adults were dissected in phosphate-buffered saline (PBS), and fixed in 2.5% glutaraldehyde in PBS overnight at 4°C. After rinsing for 30 min in PBS, the samples were post-fixed in 1% osmium tetroxide in PBS for 1 h. The samples were dehydrated in a graded series of ethanol and then infiltrated with a mixture of Epon–Araldite resin and polymerised at 60°C for 48 h. Ultrathin sections (50-70 nm thick) were cut with a Reichert ultramicrotome equipped with a diamond knife. The sections were collected with formvar-coated copper slot grids and stained with 2% aqueous uranyl acetate for 20 min in the dark and then with lead citrate for 2 min. TEM preparations were observed with a Tecnai G2 Spirit EM (FEI Eindhoven, The Netherlands) equipped with a Morada CCD camera (Olympus, Tokyo, Japan).

## Acknowledgement

This work was supported by an AFM Grant MyoNeurAlp 1 and Ligue régionale contre le Cancer. M.B. was supported by a doctoral fellowship from the AFM. J.-A.L. was supported by the FRM. Work was also supported by the ANR DIVERCIL. We acknowledge the contribution of SFR Santé Lyon-Est (UAR3453 CNRS, US7 Inserm, UCBL) facility: CIQLE (a LyMIC member). We thank J. Raff, M. Bettencourt-Dias, R. Basto, T. Megraw and G. Rogers for sharing fly stocks and reagents. Stocks obtained from the Bloomington Drosophila Stock Center (NIH P40OD018537) and the Vienna Drosophila Resource Center (VDRC) were used in this study. We thank J. Perrichet for technical assistance and fly husbandry.

## Author Contributions

VM, MB and BD designed the experiments, MB performed the majority of the experiments and analysis, VM contributed to some experiments, MGR and GC performed the electron microscopy, VM, JT and BD generated the fly stocks, VM and JAL adapted the U-ExM protocoles, VM, MB and BD wrote the manuscript.

