## Supplementary material for "Alström syndrome proteins are novel regulators of centriolar cartwheel assembly and centrosome homeostasis in *Drosophila*": Fig. S

[3] INSERM-U1314

[4] Università degli Studi di Siena, Siena, Italy

@ Equal contribution

* to whom correspondence should be addressed

Keywords: #centriole duplication #cartwheel #Ana2 #Plk4 #Alms1 #Alström syndrome

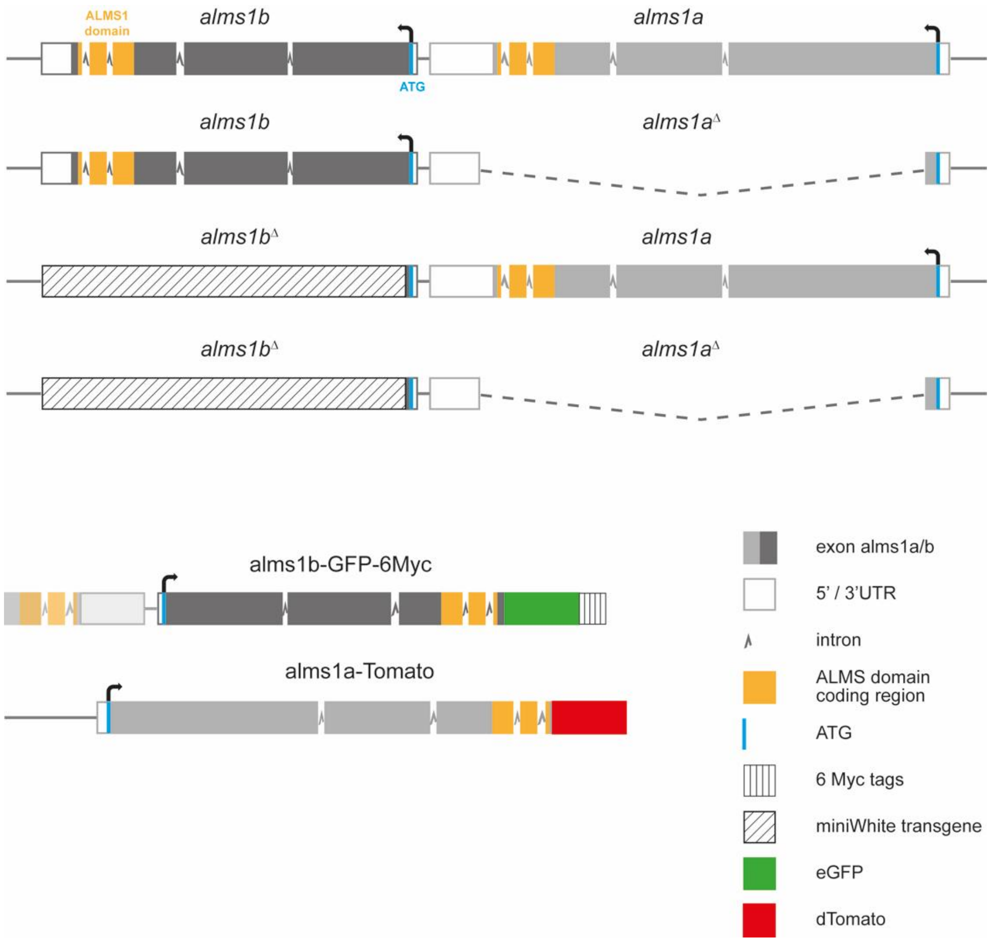


**Figure S1: Schemes of *alms1a* and *b* genetic loci and transgenic constructs**

*alms1a* (*CG12179*) and *alms1b* (*CG12184*) are localised on tandem on the minus strand of the X chromosome. The regions coding the ALMS domain are represented in orange.

*alms1a*$^{\Delta}$ was generated by NHEJ with the CRISPR/cas9 strategy. It is a 4270 nucleotides (bp) deletion between the position X: 4.635.793, after aa36 in the first exon, and the position X: 4.631.522 in *alms1a* 3'UTR. This deletion thus removes almost all *alms1a* coding sequence introducing a stop codon at position aa38 of the remaining sequence.

*alms1b*$^{\Delta}$ was generated by HDR with the CRISPR/cas9 strategy. A LoxP-miniWhite-loxP cassette was introduced between the positions and X: 4.627.368 (14nt after *alms1b* 3'UTR) and X: 4.630.821 (after aa21), it generated a deletion of 3542 bp, removing almost all *alms1b* coding sequence.

*alms1a*$^{\Delta}$, *alms1b*$^{\Delta}$ (*alms1*$^{2\Delta}$) was generated by HDR with the CRISPR/cas9 strategy on the *alms1a*$^{\Delta}$ chromosome applying the strategy used to generate *alms1b*$^{\Delta}$.

For Alms1a-Tomato transgene, the genomic sequences including *alms1a* transcribed sequence and 890 bp upstream of the transcription start site were used. dTomato coding sequence was inserted in frame after the last coding exon of *alms1a*. The transgene was inserted by targeted mutagenesis on the chromosome III at the 89E11 locus.

For Alms1b-GFP transgene, the genomic sequences including *alms1b* transcribed sequence and 1452 bp upstream of the transcription start site (which include the 3 last exons of *alms1a*) were used. A 9aa linker, the eGFP coding sequence and 6 Myc tags were inserted in frame after the last coding exon of *alms1b*. The transgene was inserted by targeted mutagenesis on the chromosome II at the 53B2 locus.

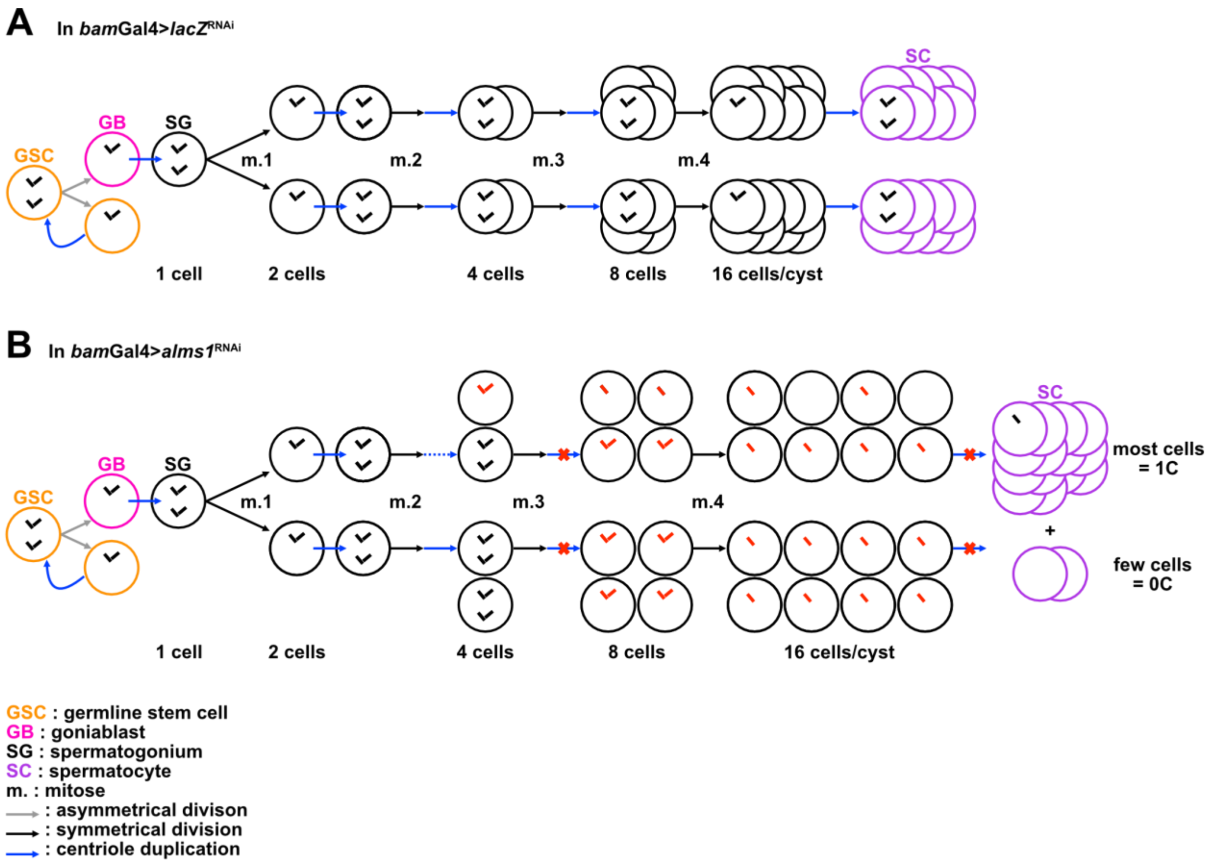


**Figure S2. Centriole duplication events during *Drosophila* spermatogenesis**

Schemes representing centriole duplication events during spermatogenesis progression in (A) *bam*-Gal4>*lacZ*$^{RNAi}$ (control condition) or (B) *bam*-Gal4>*alms1*$^{RNAi}$. Spermatogenesis starts with the asymmetric division (grey arrow) of a germline stem cell (GSC, orange) which gives rise to a GSC and a gonialblast (GB, pink) that initiates differentiation into a spermatogonium (SG, black) while it duplicates its centrioles (blue arrow). SG undergo four symmetric divisions (black arrow, mitoses m.1-4) to generate a cyst of 16 cells. Prior to each mitosis, centrioles duplicated resulting in cells with 4 centrioles arranged in two pairs. After mitose 4, SGs complete a pre-meiotic S phase and duplicate their centrioles while they differentiate into spermatocytes (SCs, purple).

*bam*-Gal4 induces the expression of the *alms1*$^{RNAi}$ (red cross) in spermatogonial cysts, and thus centriole duplication failure in some cells within 4-cells spermatogonial cysts and all cells in 8-cells cysts (centrioles in cells affected are red).

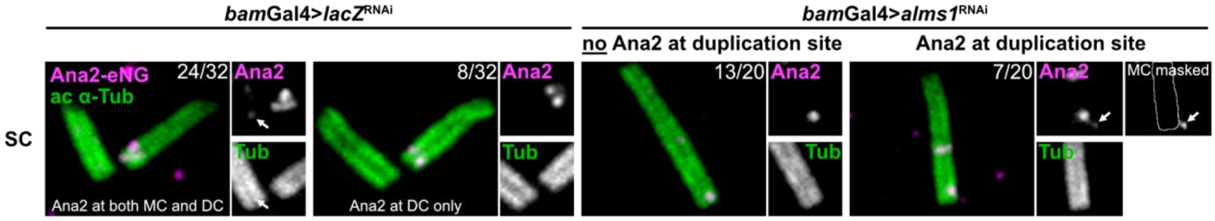


**Figure S3.**

U-ExM images of Ana2-mNeonGreen knock-in (Ana2-eNG, in magenta) in *bam*-Gal4>*lacZ*$^{RNAi}$ (control) or *bam*-Gal4>*alms1*$^{RNAi}$ in spermatocytes. Centriolar walls are revealed with acetylated $\alpha$-tubulin antibody (green). On each image, numbers indicate the occurrence of the phenotype (for SG and SC, *lacZ*$^{RNAi}$: n=71 centrioles, 3 testes; *alms1*$^{RNAi}$: n=57 centrioles, 5 testes; SG on Fig. 4). Mother centriole (MC, on the left of all images) and daughter centriole (DC, on the right). Scale bars, 1 $\mu$m.

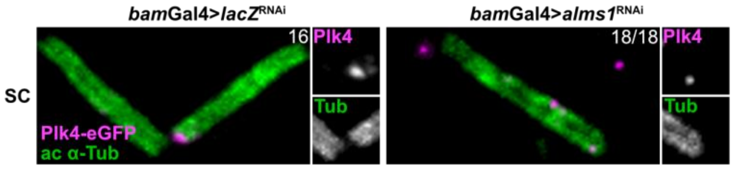


**Figure S4.**

U-ExM images of Plk4-meGFP knock-in (Plk4-eGFP, in magenta) in *bam*-Gal4>*lacZ*$^{RNAi}$ (control) or *bam*-Gal4>*alms1*$^{RNAi}$ in spermatocytes. Microtubules constituting the centriolar wall are revealed with the acetylated $\alpha$-tubulin antibody (green). On each image, numbers indicate the occurrence of the phenotype (For SG and SC, *lacZ*$^{RNAi}$: n=33 centrioles, 3 testes; *alms1*$^{RNAi}$: n=72 centrioles, 6 testes; SG on Fig. 5). Scale bars 1 $\mu$m.

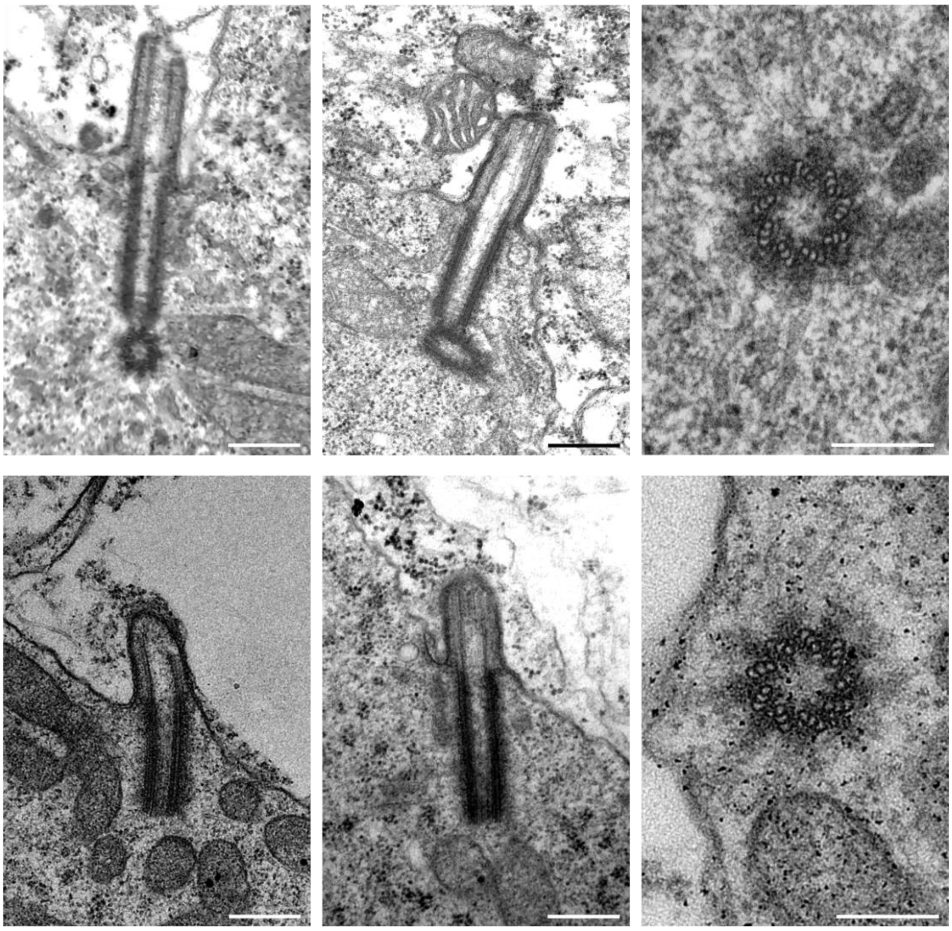


**Figure S5.** TEM images of *bam*-Gal4>*lacZ*$^{RNAi}$ (control) or *bam*-Gal4>*alms1*$^{RNAi}$ confirming the absence of procentriole formation in *alms1*$^{RNAi}$. Unduplicated centrioles show no structural defects in the centriole wall (bottom right image) and induce cilia formation (bottom left and middle images). Scale bar 5 μm on left and middle images and 200 nm on right images.
